# Integration of accessibility data from structure probing into RNA-RNA interaction prediction

**DOI:** 10.1101/359323

**Authors:** Milad Miladi, Soheila Montaseri, Rolf Backofen, Martin Raden

## Abstract

**Summary:** Experimental structure probing data has been shown to improve thermodynamics-based RNA secondary structure prediction. To this end, chemical reactivity information (as provided e.g. by SHAPE) is incorporated, which encodes whether or not individual nucleotides are involved in intra-molecular structure. Since inter-molecular RNA-RNA interactions are often confined to unpaired RNA regions, SHAPE data is even more promising to improve interaction prediction. Here we show how such experimental data can be incorporated seamlessly into accessibility-based RNA-RNA interaction prediction approaches, as implemented in IntaRNA. This is possible via the computation and use of unpaired probabilities that incorporate the structure probing information. We show that experimental SHAPE data can significantly improve RNA-RNA interaction prediction. We evaluate our approach by investigating interactions of a spliceosomal U1 snRNA transcript with its target splice sites. When SHAPE data is incorporated, known target sites are predicted with increased precision and specificity.

**Availability:** https://github.com/BackofenLab/IntaRNA

---

The function of many if not most non-coding (nc)RNA molecules is to act as platforms for inter-molecular interaction, which depends on their structure and sequence. A large number of ncRNAs regulate their target RNA molecules via base-pairing. For instance, small (s)RNAs regulate the translation of their target genes by direct RNA-RNA interactions with the respective messenger (m)RNAs [1]. To predict such interactions, regions not involved in intra-molecular base pairing have to be identified. This “free-to-interact” potential of a region, i.e. its unpaired probabilities, is computed by assessing the fraction of structures where the region is free (unpaired) within the overall structure ensemble of an RNA (see [11] for a detailed introduction). The probabilities are used by state-of-the-art prediction tools like IntaRNA [8] to account for the regions’ accessibility. While correct within their thermodynamic models, such probabilities do not incorporate all cellular constraints and dynamics that define accessible regions and thus the likelihood for interaction.

The accuracy of RNA structure prediction improves when experimental structure probing data, such as SHAPE [13], is incorporated [4]. This is done by converting the chemically sensed reactivity values to pseudo-energy terms. Pseudoenergies are combined with structure formation energies from thermodynamic models, that are used for RNA structure prediction [7, 9, 12]. As SHAPE^1^ reactivity is associated with the accessibility of nucleotides, the use of such experimental data is even more promising to improve the accuracy of RNA-RNA interaction prediction methods. For that reason, we introduce a seamless incorporation of SHAPE data into accessibility-based prediction approaches such as IntaRNA.

Recently, structure probing has been complemented by next-generation sequencing techniques to efficiently obtain transcriptome-wide reactivity information [6, 2]. This produces large data sets that demand for fast methods incorporating the probing data, which is met by our extension of IntaRNA introduced in the following.

For a given RNA-RNA interaction *I* (see supplementary material for detailed formalisms), its accessibility-based free energy is defined by *E*(*I*) = *E^hyb^*(*I*) + *ED*^1^(*I*) + *ED*^2^(*I*). Therein, *E^hyb^*(*I*) provides the hybridization energy from intermolecular base pairing while the *ED*^1,2^ terms represent the energy (penalty) needed to make the respective interacting subsequences unpaired/accessible [10]. *ED* terms are defined by unpaired probabilities *Pr^ss^* of the subsequences via *ED*(*I*) = −*RT* log(*Pr^ss^*), where *R* is the gas constant and *T* the temperature. Detailed introductions on *ED* computation are provided e.g. in [11, 14]. Computation of unpaired probabilities can be guided by SHAPE data [7]. While SHAPE-guided energy evaluations can not be compared to unconstrained energy values (due to the introduced pseudo-energy terms), unpaired probabilities are compatible, since they are reflecting the accessible structure space rather than individual structures. Thus, SHAPE-constrained 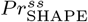 values can be directly used for *ED* and thus *E* computation while preserving comparability of the resulting energies.

Now, we show that SHAPE-guided accessibility prediction improves RNA-RNA interaction prediction. To this end, we study the probabilities of U1 small nuclear (sn)RNA interacting with pre-mRNA target sites, which is an established example of inter-molecular RNA interaction essential for RNA-splicing in eukaryotes. U1 is involved in pre-mRNA splicing by recognizing the 5’ site of introns via inter-molecular base pairing [5]. Due to the dynamics and constraints imposed by the spliceosome, it is generally challenging to avoid false positive interaction predictions, which are either predictions of U1’s recognition site with random regions of the mRNA or predicted interactions of other U1 accessible regions with the mRNA. For that reason, we used U1 as an example to show that *in vivo* probing data effectively reduces false positive predictions in RNA-RNA interaction prediction.

SHAPE data for U1 was obtained from *in vivo* DMS-seq RNA structure probing of *Arabidopsis thaliana* [3]. We selected the Ul homolog transcript bearing the largest secondary structure distance between the unconstrained and SHAPE-constrained structure prediction. The pre-mRNA sequences for 5 genes were extracted, which have been previously validated to perform U1-dependent splicing [15]. Figures 1a,b) exemplify the effect of SHAPE-constrained predictions for ACT1 mRNA. Without SHAPE constraints, the splice site is predicted to interact with various regions of U1 with high probability; see supplementary material for formalisms. In contrast, the splice site’s interaction with U1’s recognition site is dominant with high specificity in SHAPE-guided IntaRNA predictions. Furthermore, SHAPE-guided prediction has a higher precision. Among all predicted interactions of U1 with the ACT1 mRNA, the ranking of the known interaction is improved from 9 to 3 in SHAPE-guided mode. When investigating the accessibility profile of U1 (supplementary material), this mainly results from an increased SHAPE-guided unpaired probability 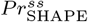 of U1’s recognition site and thus reduced *ED* penalties.

**Figure 1:**
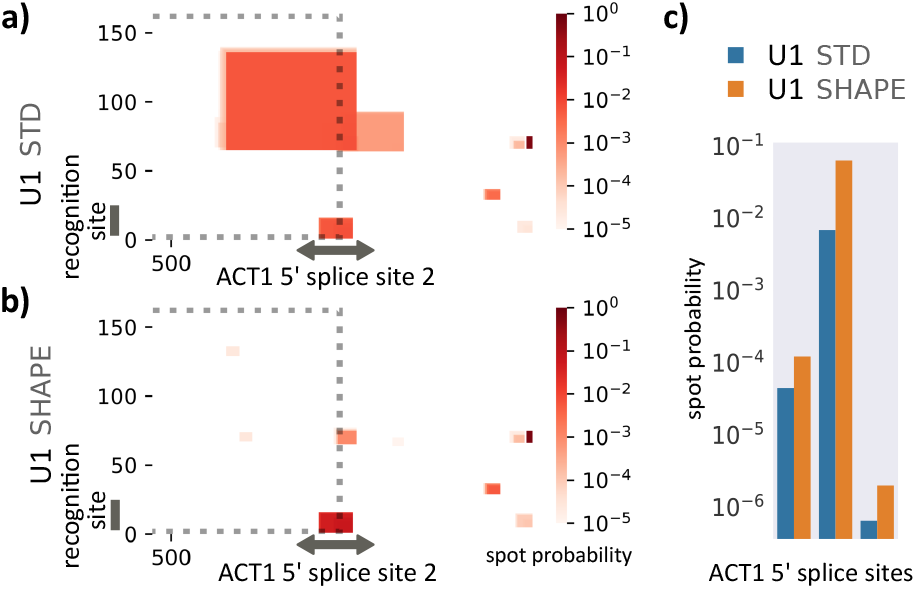
RNA-RNA interaction prediction between spliceo-somal RNA U1 with ACT1 mRNA of *Arabidopsis thaliana*. Interaction probabilities predicted between U1 (y-axis) and the region around the second intron splice site of ACT1 coding sequence mRNA using **(a)** unconstrained (STD) and **(b)** SHAPE-constrained accessibility estimates for U1. The dotted lines enclose U1 interactions with exon **2. (c)** Spot probabilities of U1 recognition site (spot index = 8) interacting with the 5’ splice sites of ACT1 (spot = 1st intron index), with SHAPE constraints (orange) and without (blue).

The interaction probability of U1’s recognition site with all three 5’ splice sites of ACT1’s coding sequence are increased when SHAPE data is incorporated (Fig. 1c). This effect results from a decreased number of false positive predictions (Fig. 1a,b). Following this trend, the probabilities of splice site recognition are improved among all the mRNAs and are on average 3.08 times higher (SHAPE/STD). Further details about data, evaluation, and analyses are provided in the supplementary material.

## Acknowledgements

We thank Dr. Ronny Lorenz for discussions on SHAPE integration.

## Supplementary material

### 1 Integrating SHAPE data into accessibility-based RNA-RNA interaction prediction

Given two RNA molecules with nucleotide sequences *S*^1^, *S*^2^ ∈ {*A*, *C*, *G*, *U*}*, we define interaction *I* between *S*^1^ and *S*^2^ as a set of inter-molecular base pairs (i.e. *I* = { (*i*,*j*) | *i* ∈ [1, |*S*^1^|] Λ *j* ∈ [1, |*S*^2^|]}), that are complementary (i.e. 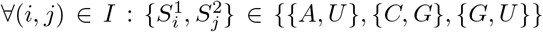) and non-crossing (i.e. ∀(*i*,*j*) ≠ (*i*′,*j*′) ∈ *I* : *i* < *i*′ → *j* > *j*′). Furthermore, any position forms at most one inter-molecular base pair (i.e. ∀(*i*,*j*), (*i*′,*j*′) : *i* = *i*′ ↔ *j* = *j*′). For any interaction *I*, the hybridization energy *E^hyb^*(*I*) can be computed using a standard Nearest-Neighbor energy model (Turner and Mathews, 2010).

The accessibility-based free energy of an interaction *I* is defined by

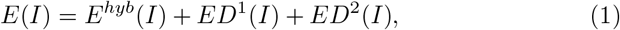

where the *ED*^1,2^(≥ 0) terms represent the energy (penalty) needed to make the respective interacting subsequences of *S*^1,2^ unpaired/accessible (Mückstein *et al*., 2006; Raden *et al*., 2018; Wright *et al*., 2018).

To compute *ED* terms, we need the left-/right-most base pair of *I* given by (*l*^1^, *r*^2^) = arg min_(*i*,*j*)∈*I*_(*i*) and (*r*^1^, *l*^2^) = arg max_(*i*,*j*)∈*I*_(*i*), respectively. Both base pairs define the interacting subsequences, i.e. 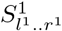 and 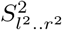. Based on that, the penalty terms are given by

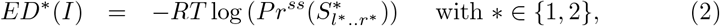

where *R* is the gas constant, *T* is the temperature, and *Pr^ss^* denotes the unpaired probability of a given subsequence, which can be efficiently computed (Bernhart *et al*., 2006).

As discussed above, SHAPE reactivity data can be incorporated into thermodynamic prediction tools via pseudo energy terms (Deigan *et al*., 2009) as has been integrated in Vienna RNA package(Lorenz *et al*., 2016). The latter enables SHAPE-guided computation of unpaired probabilities, i.e. the *Pr^ss^* terms from Eq. 2. While SHAPE-guided energy evaluations can not be compared to unconstrained energy values (due to the pseudo-energy terms), unpaired probabilities are compatible, since they are reflecting the accessible structure space rather than individual structures. Thus, SHAPE-constrained 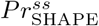 values can be directly used within the *ED* computation (Eq. 2), which provides a constrained accessibility-based interaction energy (Eq. 1) without further methodical changes. This approach is implemented in the recent version of IntaRNA e.g. available via Bioconda (Grüning *et al*., 2018).

IntaRNA interfaces all three pseudo-energy conversion methods (Zarringhalam *et al*., 2012; Deigan *et al*., 2009; Washietl *et al*., 2012) currently implemented within the Vienna RNA package. We observed best performance with Zarringhalam’s method (data not shown), which we attribute to the incorporation of reactivity on both paired and unpaired terms and was used for this study

Note, since unpaired probabilities are incorporated on negated log-scale only (compare Eq. 2), small or intermediate changes of high probability values are expected to show only minor effects in the RNA-RNA interaction prediction but should still guide or fine-tune the prediction, see Fig. 1. On the contrary, if the 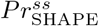 values deviate much (orders of magnitude) from *Pr^ss^* (e.g. rendering presumably unpaired regions inaccessible since they might be already blocked by other substrates or vice versa as an indirect consequence e.g. of binding) or if the probability values are very small, strong prediction effects are expected.

**Supplementary Figure 1:**
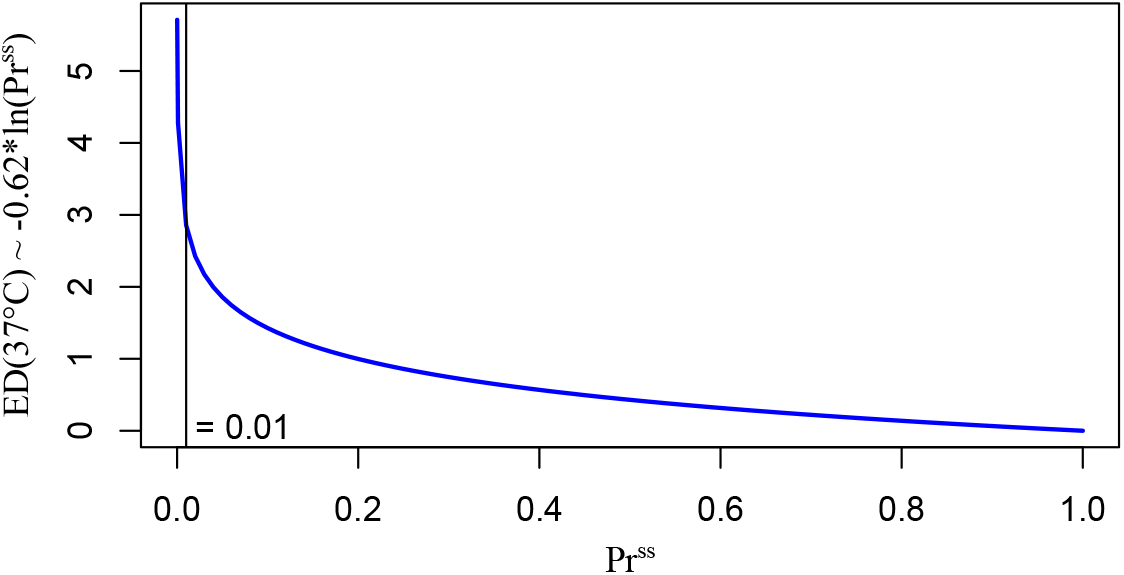
Relation of *ED* penalties and unpaired probabilities *Pr^ss^* at temperature *T* = 37°C, i.e. *RT* ~ 0.62.

### 2 Spot probabilities of RNA-RNA interaction sites

To assess the effect of SHAPE data, we define the *spot probability Pr*^spot^ of an interaction site of interest. A *spot* is defined by a pair of indices *k*, *l* for *S*^1^, *S*^2^, resp., and *Pr*^spot^(*k*, *l*) as the partition function quotient

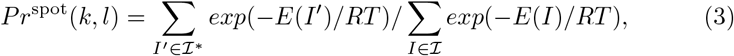

where 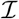 denotes the set of all possible interactions and 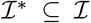 the subset of interactions that cover the spot, i.e. position *k*, *l* are within the respective interacting subsequences^1^ 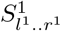 and 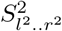 (see above).

### 3 Sequence data and evaluation scripts

All sequences used within this study as well as the used evaluation scripts are available online at

> https://github.com/BackofenLab/IntaRNA-benchmark-SHAPE

while the integration of the IntaRNA approach is available in version ≥ 2.2.0 at

> https://github.com/BackofenLab/IntaRNA

This study used IntaRNA 2.2.0, ViennaRNA v2.4.7 and pseudo energy computation for SHAPE data following Zarringhalam *et al*. (2012).

### 4 DMS-seq reactivity data extraction

We used the recommended settings from (Ding *et al*., 2015) to map the sequencing reads and compute reactivities of the annotated transcripts using the Galaxy tools Afgan *et al*. (2018) Bowtie-2 and StructureFold (Langmead and Salzberg, 2012; Tang *et al*., 2015). We selected the U1 homolog transcript bearing the largest secondary structure distance between the unconstrained and SHAPE-constrained structure prediction (using RNAfold).

### 5 U1 accessibilities with and without SHAPE reactivities

Supplementary Figure 2 depicts the accessibility of the U1 RNA in terms of position-wise unpaired probabilities computed with SHAPE data and without. Without SHAPE data, the potential of the recognition site for inter-molecular interaction is underestimated, as can be seen when comparing to the SHAPE-guided unpaired probabilities. Furthermore, the subsequent region is wrongly assumed to be accessible (while known to be blocked by intra-molecular helix formation). The probabilities correspond to the 3-mer accessibilities (RNAplfold -u 3), such that for each position i the probability that 3-mer [i-2, i] is unpaired.

This can also be seen from the accessibility-annotated structure plots in Supplementary Fig. 3. The accessibility (”unpairedness”) of the recognition site is much better reflected by SHAPE-guided terms compared to standard computations.

**Supplementary Figure 2:**
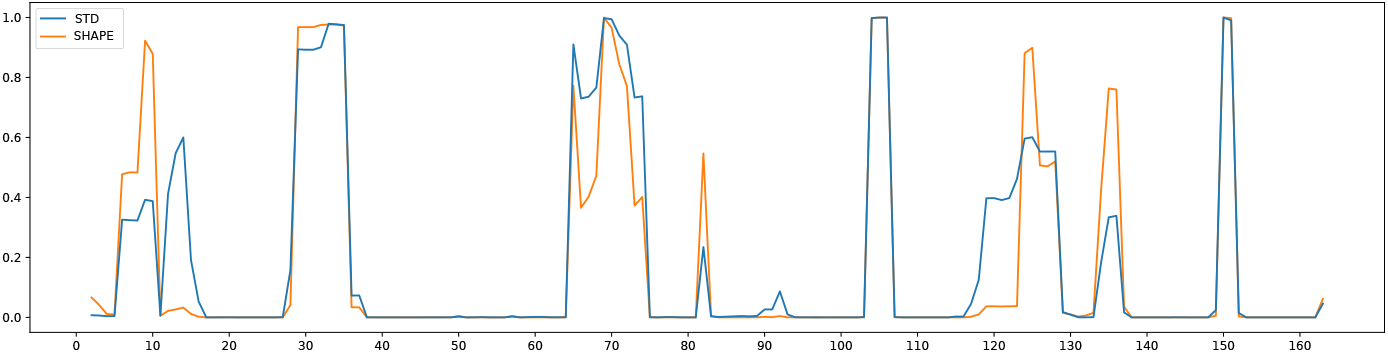
Position-wise accessibility (y-axis) of U1 with SHAPE data (orange) and without (blue, STD).

**Supplementary Figure 3:**
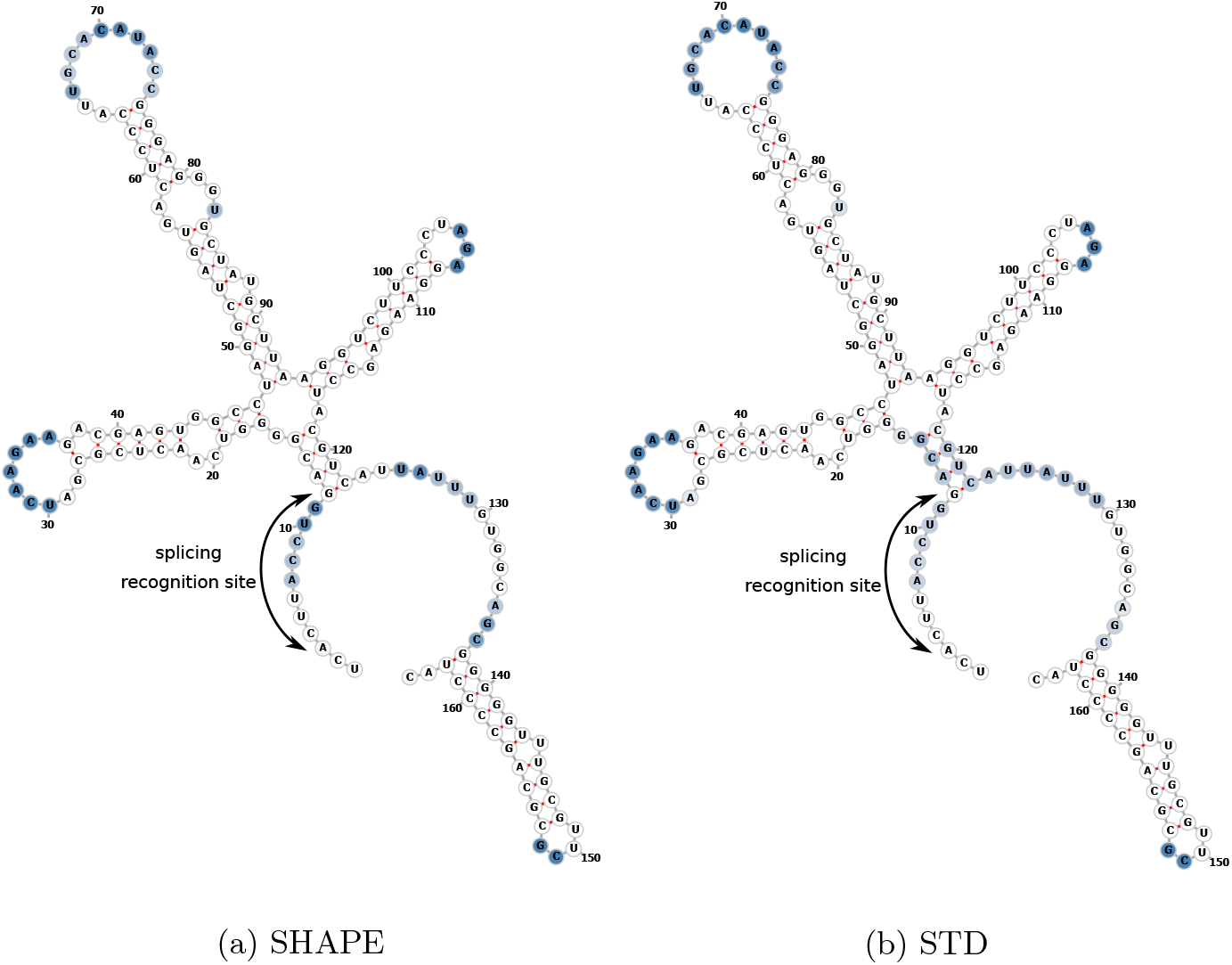
Accessibilities mapped to the U1 secondary structure and color-coded, more accessible positions have darker colors. Visualization using Forna web-server (Kerpedjiev *et al*., 2015)

### 6 Pre-mRNA U1 interaction predictions details

In the following, we provide the interaction prediction heatmap for all studied mRNAs. Index combinations that mark interactions of U1 with coding exons are enclosed by gray dotted boxes. The coloring represents the respective spot probabilities. Each heatmap is complemented with a visualization of the spot probabilities of U1’s recognition site interacting with each CDS 5’-splice site using SHAPE data and without.

Predicted interaction sites are drawn in colored boxes where darker boxes relate to higher interaction (spot) probabilities. The x-axis corresponds to pre-mRNA indices while the y-axis represents positions of U1. For the latter, the U1 recognition site for intronic 5’ splice sites is at position 4-11. Thus, interactions of that site correspond to the bottom of the graph. The top graph represents interaction sites without SHAPE constraints while the bottom graph depicts the altered prediction when U1 SHAPE constraints are considered within IntaRNA’s accessibility computation.

The left barplot shows the predicted interaction probabilities of all intronic 5’ splice sites with the recognition site of U1 (called a spot). Blue bars represent the unconstrained probabilities while orange bars depict the probabilities when U1 SHAPE constraints are used. The right bar plot provides the ratio of both probabilities for each interaction site (spot). Please note, representations of multiple spots are in log-scaling to enable depiction.

**Supplementary Figure 4:**
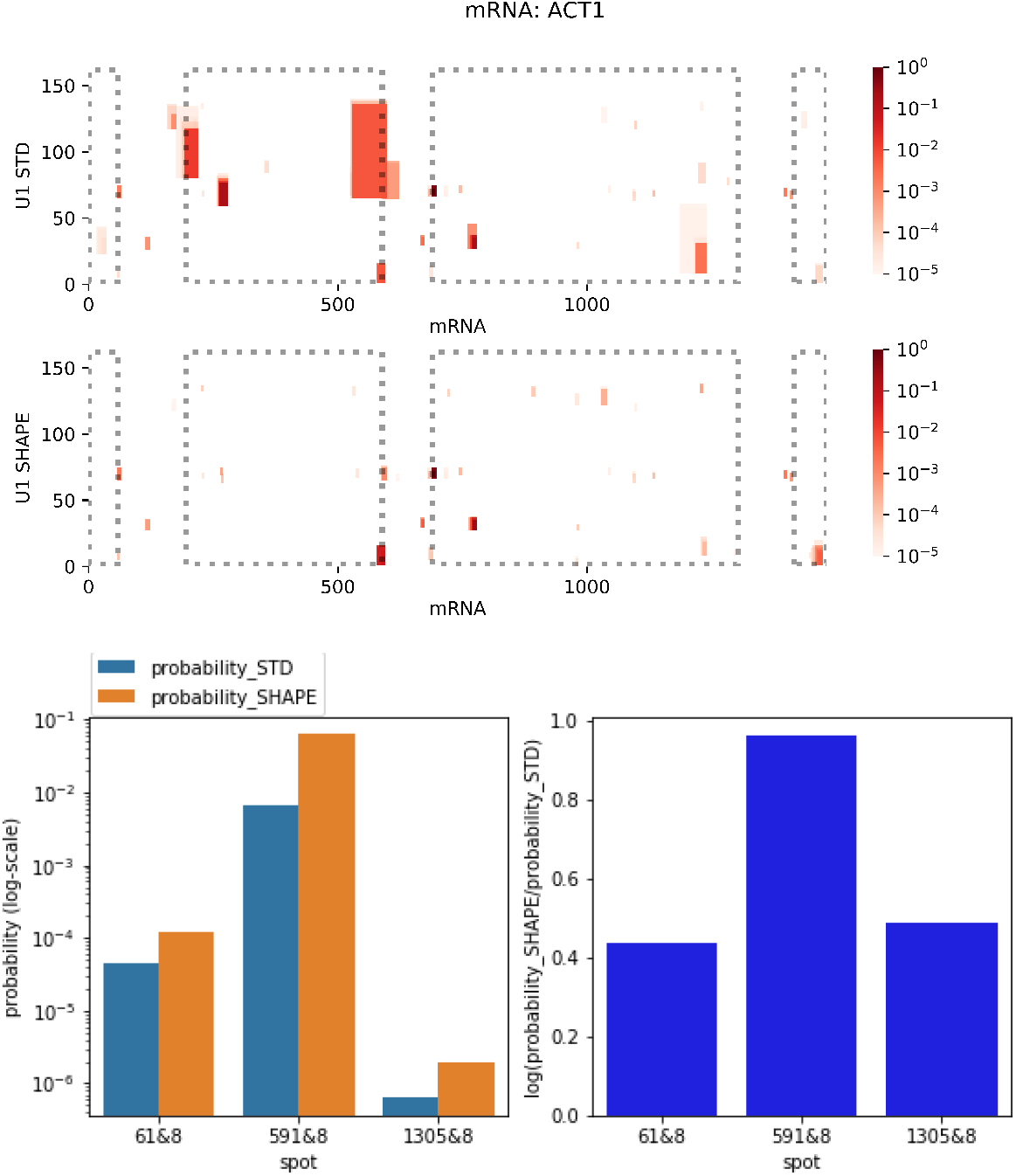
U1-ACT1 interaction prediction with and without U1 probing data. Note, bar plots are in log_10_-scale.

**Supplementary Figure 5:**
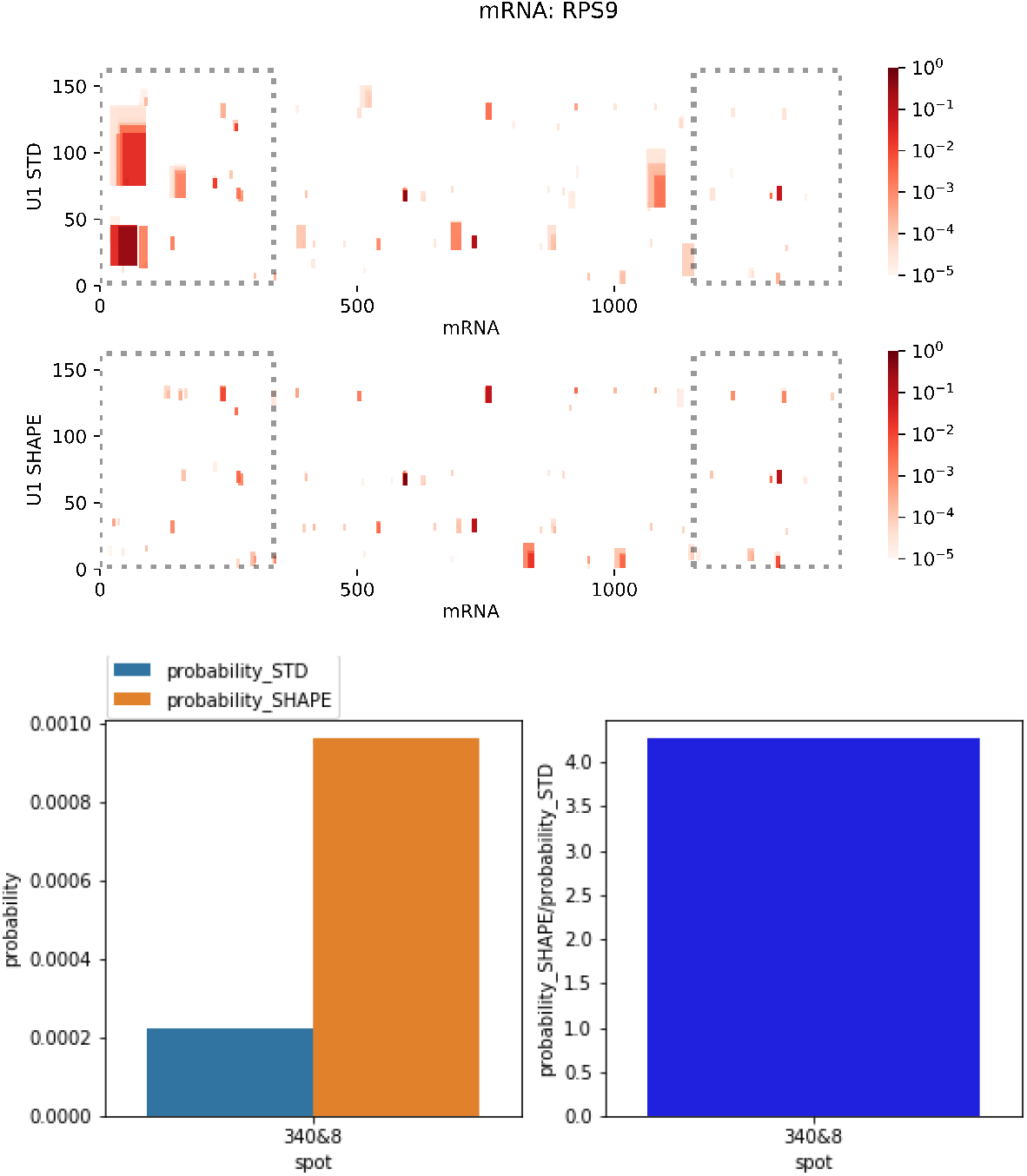
U1-RPS9 interaction prediction with and without U1 probing data

**Supplementary Figure 6:**
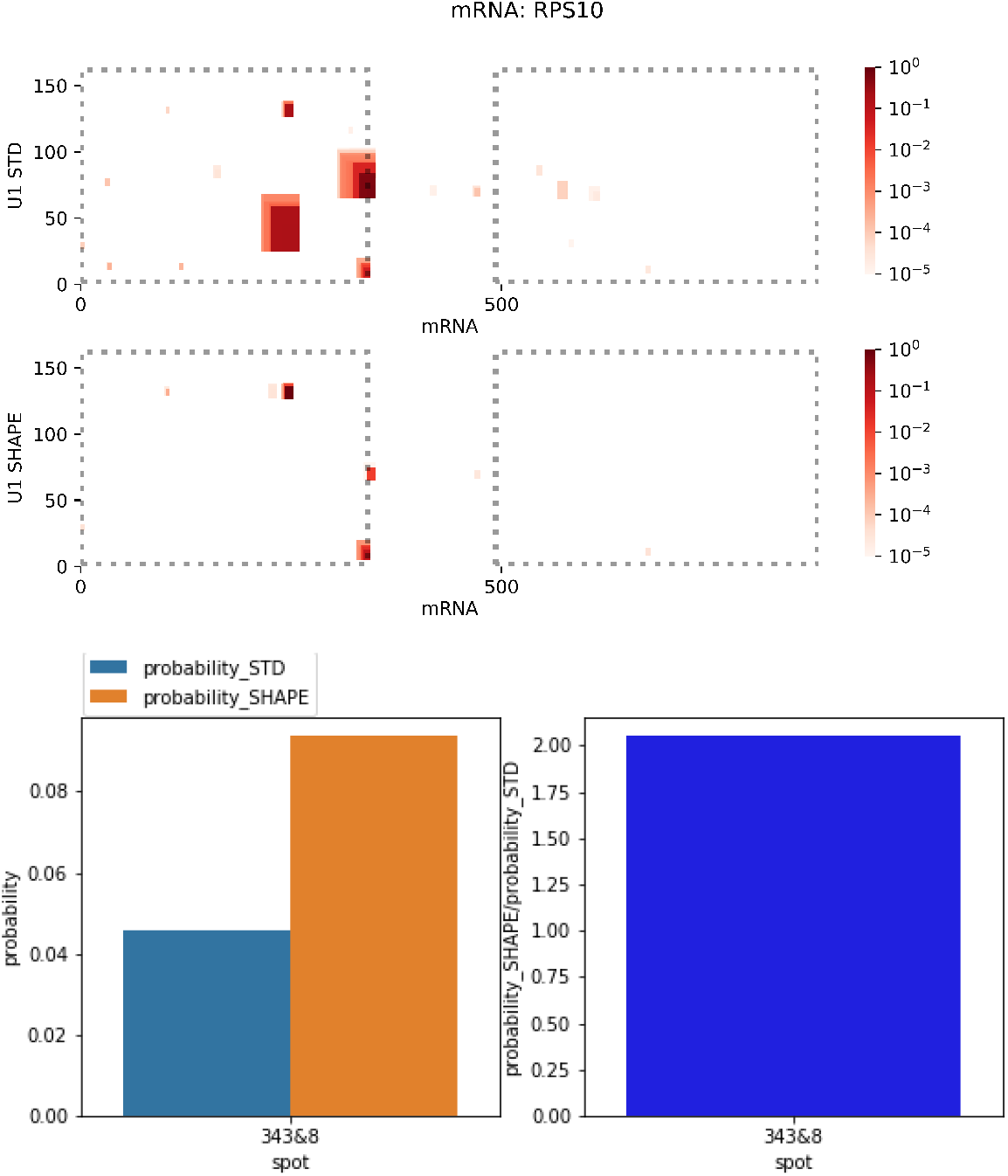
U1-RPS10 interaction prediction with and without U1 probing data

**Supplementary Figure 7:**
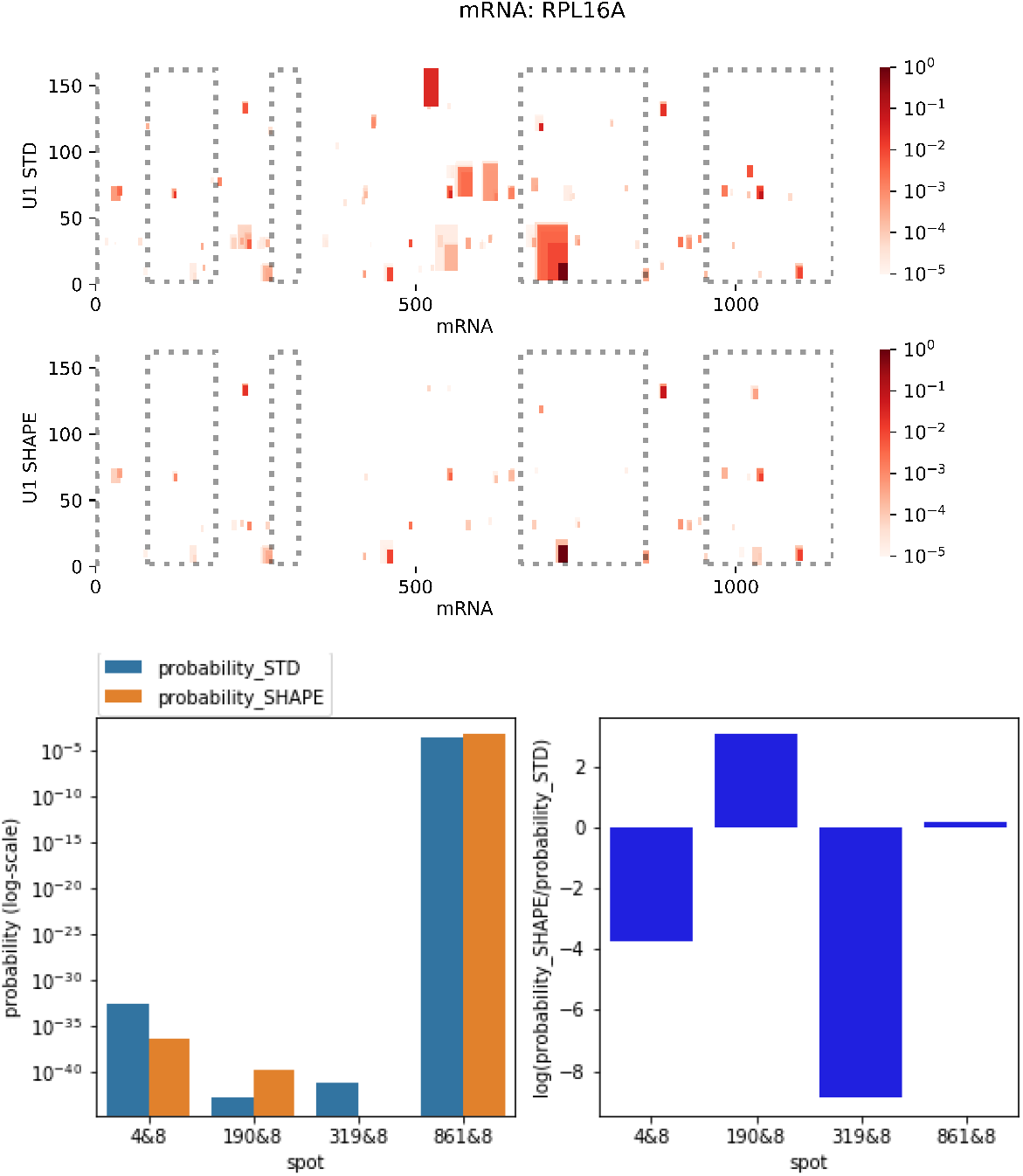
U1-RPL16A interaction prediction with and without U1 probing data. Note, bar plots are in log_10_-scale.

**Supplementary Figure 8:**
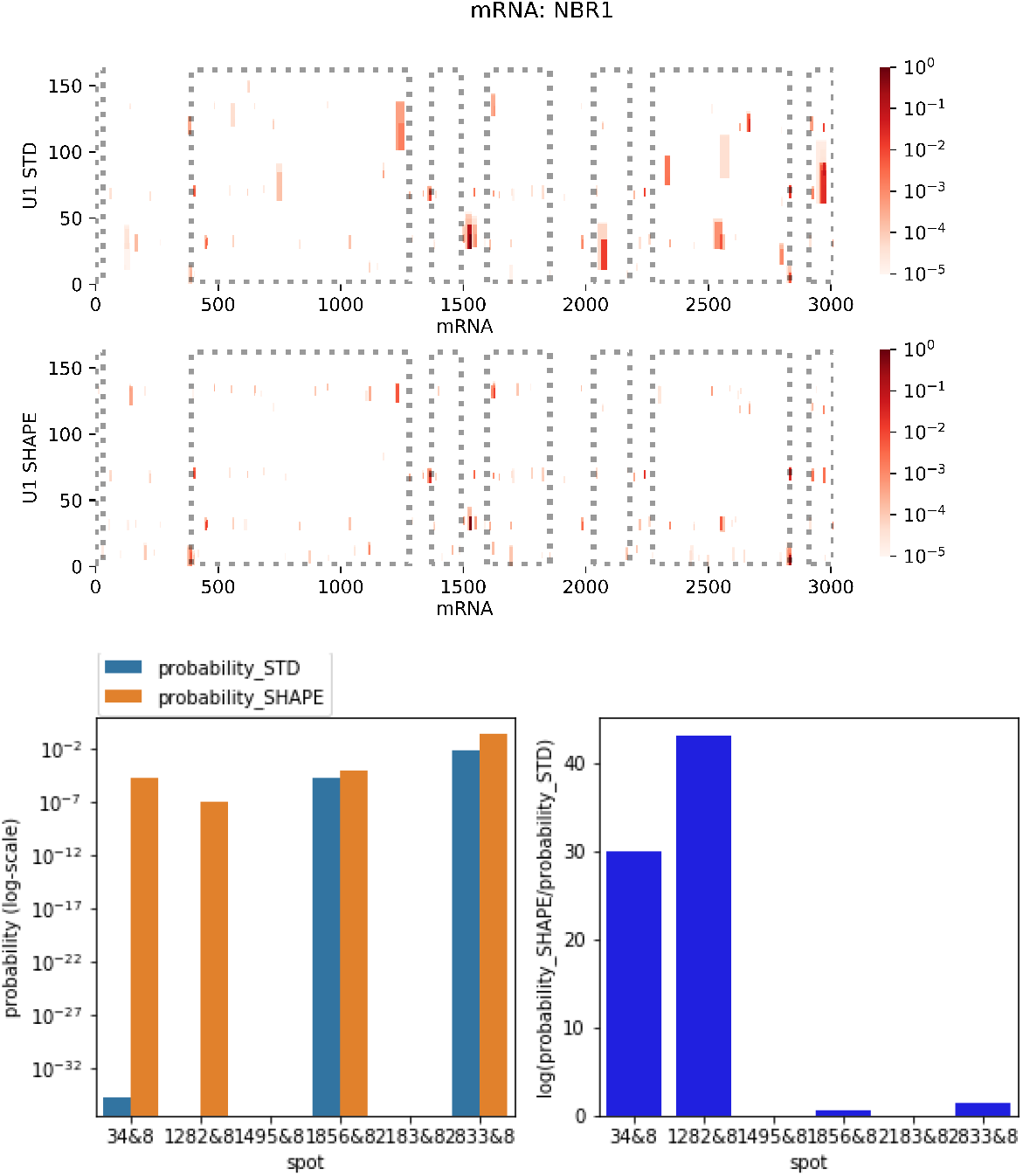
U1-NBR1 interaction prediction with and without U1 probing data. Note, bar plots are in log_10_-scale.

## Footnotes

1 For simplicity, SHAPE refers to any structure probing experiment (e.g. SHAPE, DMS).

1 Note, interactions 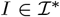 covering a spot at *k*,*l* do not necessarily contain the base pair (*k*,*l*), i.e. *k*, *l* or both can be unpaired.

